# Survival and Inactivation by Advanced Oxidative Process of Foodborne Viruses in Model Low-Moisture Foods

**DOI:** 10.1101/2020.03.05.980078

**Authors:** Neda Nasheri, Jennifer Harlow, Angela Chen, Forest Dussault, Nathalie Corneau, Sabah Bidawid

**Affiliations:** National Food Virology Reference Centre, Bureau of Microbial Hazards, Food Directorate, Health Canada 251 Sir Frederick Banting Driveway, Ottawa, ON, K1A 0K9, CANADA; Biostatistics and Modeling Division, Bureau of Food Surveillance and Science Integration, Food Directorate, Health Canada Ottawa, ON, Canada

**Keywords:** Hepatitis A virus, norovirus surrogates, low moisture foods, bead-based assay, whole genome sequencing, advanced oxidative process

## Abstract

Enteric viruses, such as human norovirus (NoV) and hepatitis A virus (HAV), are the major causes of foodborne illnesses worldwide. These viruses are shed in high numbers, have low infectious dose, and may remain infectious for weeks in the environment and food. While numerous viral survival studies have been conducted in fresh fruits and produce, limited information is available regarding viral survival and transmission in low moisture foods (LMF). LMFs are generally considered as ready-to-eat products, which undergo no or minimal pathogen reduction steps. However, numerous foodborne outbreaks associated with LMFs have been reported in recent years. The objective of this study was to examine the survival of foodborne viruses in LMFs during long-term storage at ambient temperature and to evaluate the efficacy of advanced oxidative process (AOP) treatment in the inactivation of these viruses. For this purpose, select LMFs such as pistachios, chocolate, and cereal were inoculated with HAV and the norovirus surrogates, murine norovirus (MNV) and feline calicivirus (FCV), then viral survival on these food matrices was measured over a four-week incubation at ambient temperature, by both plaque assay and droplet-digital RT-PCR (ddRT-PCR). We observed an approximately 0.5 log reduction in viral genome copies, and 1 log reduction in viral infectivity for all three tested viruses following storage of select inoculated LMFs for 4 weeks. Therefore, the present study shows that foodborne viruses can persist for long-time in LMFs. Next, we examined the inactivation efficacy of AOP treatment, which combines UV-C, ozone, and hydrogen peroxide vapor, and observed that while approximately 100% inactivation can be achieved for FCV, MNV, and HAV in chocolate, the inactivation efficiency diminishes to approximately 90% in pistachios and 70% in cereal. AOP treatment could therefore be a good candidate for the elimination of foodborne viruses from certain LMFs depending on the food matrix and surface of treatment.

**Importance:** Low moisture foods have been increasingly recognized as important vehicles of foodborne pathogens. In the present study, we demonstrated that foodborne viruses remain infectious during long-term storage on select low moisture foods. In addition, we evaluated the efficacy of an advanced oxidative process in the inactivation of foodborne viruses in low moisture foods. This research will help increase the safety of low moisture foods and reduce the number of foodborne illnesses due to contaminated products.

## Introduction

Gastroenteritis outbreaks and illnesses due to the consumption of contaminated low moisture foods (LMFs) are recently emerging as food safety burden in developed countries, with norovirus and hepatitis A virus being important causes of LMF-associated outbreaks (1–5).

Due to their low water activity (a_w_), LMFs are considered less susceptible to the growth of foodborne pathogens (6). For this reason, minimal intervention strategies are implemented during the production of LMFs (6). However, foodborne viruses are relatively resistant to various environmental factors (e.g., pH, temperature, and a_w_), and viral persistence in various environments such as food surfaces and soil, has been demonstrated (7–9). The environmental stability of foodborne viruses together with their low infectious dose and resistance to conventional antimicrobial treatments, could lead to their persistence in LMFs and cause serious implications on public health.

Little is known about the survival and inactivation of viruses in LMFs, while recent outbreaks indicate that LMFs can be important vectors for foodborne viruses. In our previous study, we have developed a magnetic bead-based assay for the isolation of HAV and norovirus surrogates, feline calicivirus (FCV) and murine norovirus (MNV), from LMFs and compared its efficacy against the adapted ISO 15216-1 method (10). We have demonstrated that while both methods can be employed to recover viruses from LMFs, each method has its own advantages and disadvantages (10, 11). Thus, both methods were applied in the present study to investigate viral persistence and infectivity during four-week storage in LMFs. Three popular LMFs that are usually consumed without further processing (i.e. chocolate, pistachios, and cereals), were selected for this study. Viral survival on each matrix was examined by plaque assay as well as droplet-digital RT-PCR (ddRT-PCR).

We have also conducted next generation sequencing (NGS) to investigate genomic alterations that occur in viral quasispecies during the incubation on LMFs, and we characterized the genomic differences between the viruses isolated after long-term incubation in select LMFs and the viruses that were extracted at time zero (T_0_).

Since it is crucial to find an effective tool to mitigate enteric viruses from ready-to-eat LMFs, we explored the efficiency of advanced oxidative process (AOP), which combines UV-C radiation with ozone and hydrogen peroxide vapor treatments for the inactivation of pathogens in ready-to-eat foods. UV-C treatment has been used with relative success to eliminate foodborne viruses in fresh produce and food-contact surfaces (reviewed in (12, 13)). The viricidal efficacy of ozone against foodborne viruses is variable and depends on several factors including the matrix and the ozone concentration (dose) (14). It is believed that ozone treatment affects the viral capsid and genome, however, its inactivation mechanism is poorly understood (15). Hydrogen peroxide (H_2_O_2_) acts as an oxidant against bacteria and viruses by producing hydroxyl-free radicals, which affect viral capsid and genome. Moreover, H_2_O_2_ breaks down to water and oxygen and thus constitutes a low environmental risk (16, 17). The efficacy of H_2_O_2_ treatment against HAV and surrogates of norovirus has been shown to be matrix-dependent (reviewed in (12, 13)). In this study, we evaluated the combinational effect of these inactivation strategies against foodborne viruses in select LMFs by cell culture based infectivity assays as well as ddRT-PCR assay.

## Materials and Methods

### Cells and viruses

Crandell Rees feline kidney (CrFK) cells were obtained from ATCC (# CCL-94) and maintained as previously described (18). Feline Calicivirus (FCV) (ATCC # VR-782) was used to inoculate LMFs.

Murine BV-2 cells (kindly provided by Dr. Wobus, University of Michigan, Ann Arbor) were grown in Dulbecco’s modified Eagle’s medium (Invitrogen, Burlington, Ontario, Canada) supplemented with 10% fetal bovine serum, 0.1 mg/ml L-glutamine, and 0.1 mg penicillin-streptomycin. The cells were incubated at 37°C with 5% CO_2_ and maintained as previously described (19). MNV (kindly provided by Dr. Virgin, Washington University School of Medicine, St. Louis, MO) was used to inoculate LMFs.

Hepatitis A virus strain HM-175 (ATCC # VR-1402) and seed cultures of fetal rhesus monkey kidney cells (FRhK-4) were kindly provided by Dr. S. A. Sattar (University of Ottawa). FRhK-4 cells were grown, maintained and prepared as described previously (20).

### Plaque assay

Virus titres were quantified by plaque assay in 12-well plates (Millipore Sigma, Cat# Z707775). For the quantification of FCV, MNV, and HAV, the cell lines CrFK, BV-2, and FRhk-4, respectively, were grown, maintained, and infected with the different low moisture food samples. The cell line monolayers were grown at 37°C overnight, and each of three wells was inoculated with 100 µl of each sample (pistachios, chocolate or cornflakes) and sample dilution. Samples were diluted using 1 × EBSS (Thermofisher, Cat# 24010-043). Samples were adsorbed for 60 min at 37°C, with gentle rocking of the plates every 10 min to evenly distribute the sample on the cell monolayer. A 2 ml mixture of agarose-media was overlaid onto monolayers and the plates were incubated at 37°C with 5% CO_2_ for 2 days (FCV and MNV) or 8 days (HAV). The monolayers were fixed with 3.7% paraformaldehyde (Millipore Sigma, Cat# F1635) for a minimum of 4 h. The monolayer was stained with 0.1% crystal violet and plaques were counted manually and converted to PFU/ml.

### Inoculation

Chocolate liqueur (Cargill) was weighed in 4 g amounts into wells of a 6-well plate. Chocolate was melted at 43°C in a bead bath. The chocolate solidified and was stored at room temperature until samples were ready for virus inoculation. Chocolate samples were inoculated with 100 µl of 5 × 10^6^ PFU/ml of FCV (6.2 × 10^5^ genome copies/µl), 8 × 10^7^ PFU/ml of MNV (1.05 × 10^6^ genome copies/µl), or 1 × 10^5^ PFU/ml of HAV (1.2 × 10^5^ genome copies/µl) by pipetting small volumes at a time into each well to ensure even distribution of the virus. The inoculated chocolate was dried at room temperature (RT) in a biosafety level 2 cabinet (BSC) for 1 h. Samples were inoculated in triplicate (3 wells of 4 g of chocolate per time point) in two sets. The samples at T_0_ were analysed immediately and the rest of the samples were incubated at RT for 24 h, 1 week, 2 weeks, 3 weeks and 4 weeks for survival assay prior to extraction.

Twenty-five grams of unsalted dry-roasted, shelled pistachios from American Pistachio Growers (APG) were weighed in petri dishes in two sets, each in triplicate. The pistachio samples were inoculated with 100 µl of 5 × 10^6^ PFU/ml of FCV (6.2 × 10^5^ genome copies/µl), 8 × 10^7^ PFU/ml of MNV (1.05 × 10^6^ genome copies/µl), or 1 × 10^5^ PFU/ml of HAV (1.2 × 10^5^ genome copies/µl). Each pistachio was inoculated with a small droplet of virus stock, until all 25 g of the pistachios were covered with at least one virus droplet, and dried in a BSC at room temperature for 1 h. The samples at T_0_ were analysed immediately and the rest of the samples were incubated at RT for 24 h, 1 week, 2 weeks, 3 weeks and 4 weeks for survival assay prior to extraction.

Cornflakes (Kellogg’s) were weighed in 25 g amounts in petri dishes in two sets, each in triplicate. A 100 µl volume of 5 × 10^6^ PFU/ml of FCV (6.2 × 10^5^ genome copies/µl), 8 × 10^7^ PFU/ml of MNV(1.05 × 10^6^ genome copies/µl), or 1 × 10^6^ PFU/ml of HAV (1.2 × 10^5^ genome copies/µl) was used to inoculate the cornflakes and dried in a BSC at room temperature for 30 min. The samples belong to T_0_ were analysed immediately and the rest of the samples were incubated at RT for 24 h, 1 week, 2 weeks, 3 weeks and 4 weeks for survival assay prior to extraction.

### ISO 15216-1:2017 method (21)

To recover FCV, MNV, or HAV, from the inoculated chocolate, the entire surface of the chocolate was swabbed five times with a sterile cotton swab pre-moistened with 1 × phosphate-buffered saline (PBS, pH 7.2). The swab was then immersed in 500 µl PBS for virus quantification by plaque assay or lysis (AVL) buffer for viral RNA extraction using Qiagen’s QIAmp Viral RNA mini kit according to manufacturer’s instructions.

To recover viruses from inoculated pistachios, the 25 g of pistachios were added to mesh filter bags (VWR, #11216-904, PA) and 40 mL of TBGE buffer (100 mM Tris pH 9.5, 50 mM glycine, 1% wt/vol beef extract) was added to each sample. The sample was mixed and incubated at room temperature, on a rocking plate for 20 min. The eluate was transferred to a 50 ml centrifuge tube and centrifuged at 10,000 × g for 30 min at 4°C. The supernatant was decanted into a new tube and the pH was adjusted to 7±0.5 with HCl. 5 × polyethylene glycol 6000 (500g/l PEG, Sigma)/NaCl (1.5mol/l Fisher) of ¼ volumes of the weight of each sample was added to each tube, followed by incubation on ice, on a rocking plate for 1 h. Samples were centrifuged at 10,000 × g for 30 min at 4°C. The supernatant was discarded and the pellet was resuspended in 500 µl of PBS and stored at −80°C for further analysis by plaque assay or ddRT-PCR.

To recover viruses from inoculated cornflakes, the 25 g of cornflakes were added to mesh filter bags and 175 ml of TGBE buffer were added to each sample. The sample was mixed and incubated at room temperature, on a rocking plate for 20 min. The eluate was transferred to a 50 mL centrifuge tube and centrifuged at 10,000 × g for 30 min at 4°C. The supernatant was decanted into a new tube and the pH was adjusted to 7±0.5 with HCl. Samples were frozen at − 80°C overnight. Samples were thawed the next day, and 5 × PEG/NaCl of ¼ volumes of the weight of each sample was added to each tube, followed by incubation on ice on a rocking plate for 1 h. Samples were centrifuged at 10,000 × g for 30 min at 4°C. The supernatant was decanted and discarded and the pellet was resuspended in 1000 µl – 2000 µl PBS depending on the size of the pellet for further analysis by plaque assay or ddRT-PCR.

Following viral recovery from inoculated LMFs, viral RNA extraction was performed using QIAmp Viral RNA mini kit according to manufacturer’s instructions.

### FCV and MNV recovery using bead-based method

A 1:10 bead to sample ratio was used for all the bead-based extractions (100 µl of beads for 1ml of eluate). Porcine gastric-coated magnetic beads (PGM-MB) were prepared as described previously (11, 22). To recover FCV or MNV from inoculated chocolate using the bead-based method, the surface of the chocolate was swabbed with a pre-moistened cotton swab five times and the virus was released in 800 µl PBS and extracted using 80 µl PGM-MB beads.

To recover FCV or MNV from inoculated pistachios using the PGM-MB method, 25 g of virus inoculated pistachios were placed in mesh filter bags with 40 ml PBS. The samples were incubated at room temperature on a rocking plate for 20 min. The eluate was transferred to a 50 ml centrifuge tube and 1 ml of eluate was used for extraction using 100 µl PGM-MBs.

To recover FCV or MNV from inoculated cornflakes using the PGM-MB method, 25 g of virus-inoculated cornflakes were placed in mesh filter bags with 175 ml of PBS. The samples were incubated at room temperature, on a rocking plate for 20 min. The eluate was transferred to a 50 mL centrifuge tube and 1 ml of eluate was used for extraction using 100 µl PGM-MBs.

For RNA extraction, Samples were incubated with PGM beads for 30 min on a rotary shaker. The beads were washed with PBS three times and resuspended in 50 µl ultrapure water, heated at 100°C for 10 min and then quickly chilled on ice. RNA was stored at −80°C.

### Virus recovery using anti-HAV conjugated beads

HAV-Ab coupled beads were prepared using the magnetic DynaBeads Antibody Coupling Kit (Thermofisher, Cat# 14311D) according to manufacturer’s instructions. Briefly, 10 mg of M-270 Epoxy Dynabeads were weighed and added to 1 ml C1 buffer. Samples were incubated on a rotary shaker for 5 min. In a separate tube, 450 µl of C1 and 50 µl of HAV antibody (Creative Diagnostics, Cat# DPA0224) was combined and added to the beads. A 500 µl volume of C2 was added and incubated on a roller at 37°C for 16-24 h. The beads were washed once with HB and LB, and washed twice with SB. The solution was incubated on a roller for 15 min at room temperature. The supernatant was removed, and the beads were resuspended in 1 mL SB and stored at 4°C.

To recover HAV from inoculated chocolate using the bead-based method, the surface of the chocolate was swabbed with a pre-moistened cotton swab five times and the virus was released in 800 µl PBS and extracted using 80 µL of anti-HAV conjugated beads.

To recover HAV from inoculated pistachios using the bead method, 25 g of virus-inoculated pistachios were placed in mesh filter bags with 40 ml PBS. The samples were incubated at room temperature on a rocking plate for 20 min. The eluate was transferred to a 50 ml centrifuge tube and 1 mL of eluate was used for RNA extraction using 100 µL anti-HAV conjugated beads.

To recover HAV from inoculated cornflakes, 25 g of virus-inoculated cornflakes were placed in mesh filter bags with 175 mL PBS. The samples were incubated at room temperature on a rocking plate for 20 min. The eluate was transferred to a 50 ml centrifuge tube and 1 mL of eluate was was used for RNA extraction using 100 µL anti-HAV conjugated beads.

The anti-HAV conjugated beads were washed with PBS + 0.1% BSA (Millipore Sigma, Cat# A2153). Eluates containing viruses were added to the beads and incubated at room temperature for 1 h on a rotary shaker. The beads were washed with PBS four times, and resuspended in 50 µL of ultrapure water, heated at 100°C for 10 min and then quickly chilled on ice. RNA was stored at −80°C.

### ddRT-PCR

FCV, MNV or HAV RNA recovered by the ISO 15216-1 and PGM-MB or HAV antibody methods was quantified using the One-Step dd-RT-PCR Advanced Kit for Probes (Bio-Rad Laboratories, Ltd.) according to manufacturer’s instructions and as described previously (23, 24). Primers and probes used to quantify FCV, MNV and HAV are listed in Table 1.

**Table 1.**
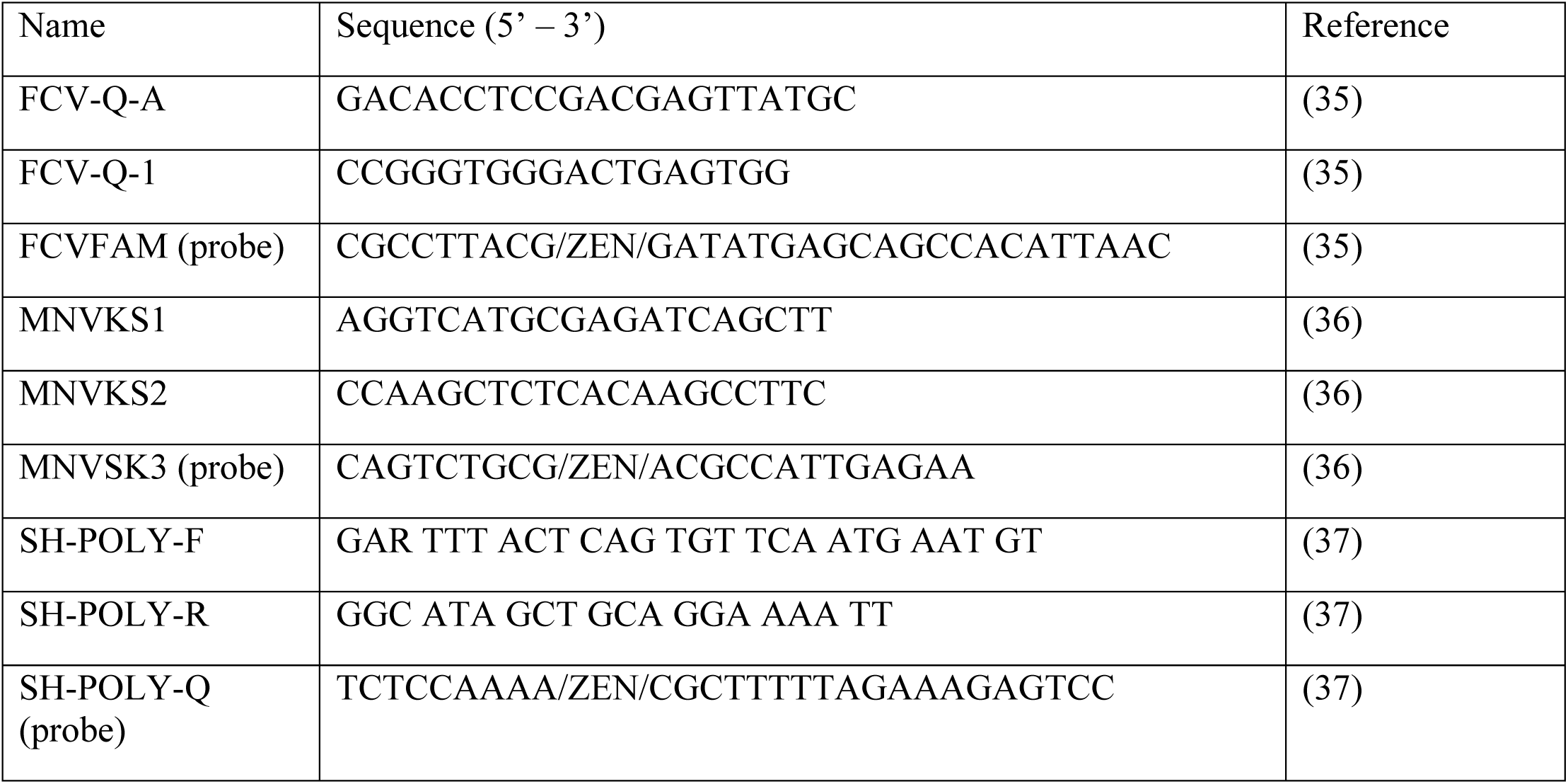
List of primers and probes used in this study. The probes were labelled with Black quencher and FAM reporter dye for FCV and MNV, and HEX reporter dye for HAV

The probes for the detection and quantification of the viruses in this study were made with the Iowa Black quencher (Integrated DNA technologies), as this quencher produced the least background. The FAM reporter dye was used for probes to detect FCV and MNV, and the HEX reporter dye was used for HAV. All probes were made with an internal ZEN quencher to reduce background noise (Integrated DNA Technologies).

The thermocycling conditions were: 50°C for 60 min, 95°C for 10 min, 40 cycles of 95°C for 30s (Ramp = 2°C/sec) with an annealing temperature of 55°C for FCV, and 53°C for MNV and HAV, for 1 min (Ramp = 2°C/sec), and 98°C for 10 min. ddRT-PCR results were analyzed using the QX200™ Droplet Digital system (Bio-Rad Laboratories Ltd.).

### Statistical analysis

Statistical analysis was performed using Microsoft Excel 2016 to determine significant differences between the recovery rates obtained by both methods using the paired t-test.

### Whole genome sequence analysis (WGS)

FCV, MNV, and HAV RNA isolated from pistachios, chocolate and cornflakes using the ISO15216 method from two time points, T_0_ and W4, were selected for WGS analysis. Ethanol precipitation of RNA was performed prior to proceeding to TruSeq Stranded mRNA (Illumina, San Diego, California, USA) sample preparation according to the manufacturer’s instructions and as described before (25).

### Bioinformatic analysis and sequence assembly

To generate consensus sequences for the samples, first MiSeq short reads were aligned against their respective reference genomes using BBMap v38.54 (26) to generate a sorted BAM file. Next, genomic variants were generated with output from the Samtools (27) mpileup utility piped into the BCFTools (27, 28) multiallelic call utility. This resulted in a variant file, which was used with the sample’s respective reference genome and processed with the BCFTools consensus module, producing a final consensus genomic sequence. Finally, coverage statistics were generated with the BEDtools (29) genomecov utility.

### Inactivation by Advanced Oxidation Process (AOP)

Twenty-five gram samples of each low moisture food were inoculated with 100 µl of 5 × 10^6^ PFU/ml of FCV, 8 ×10^7^ PFU/ml of MNV, or 1 × 10^5^ PFU/ml of HAV. In triplicate, each food was exposed to 0 seconds (untreated), 30 seconds and 1 minute of UV and hydrogen peroxide treatment. UV and hydrogen peroxide treatment was applied to the samples using an AOP device (Clean Works^©^, Beamsville, ON Canada), which is an instrument consisting of a conveyor belt that moves through a treatment chamber with spray nozzles that release hydrogen peroxide (6%) at a flow rate of 50ml/min, 4 UV-C bulbs (254 λ, 82 ϕ/watts each), and 4 ozone generating bulbs (187 λ, 2.3 mg/hour ozone each).

For 30 seconds of treatment, the samples (pistachios and cornflakes in petri dishes, and chocolate in 6-well plates) were placed in a metal container on the conveyor belt and passed though the treatment chamber for 30 seconds (the manufacturer’s recommendation). For 60 seconds of treatment, the samples were placed in metal containers on the conveyor belt and held for an additional 30 seconds by attaching masking tape to the metal dish. Viruses were recovered from inoculated LMFs using the ISO-15216-1 method. Quantification of viral RNA was done by ddRT-PCR and the number infectious viruses was quantified by plaque assay as described above.

## Results

### Virus survival in low moisture foods

We examined the survival of FCV, MNV and HAV in LMF by inoculation of pistachios, chocolate, cereals, and storage at room temperature (RT) for four weeks. The survival of the viruses on chocolate and pistachios was determined using both the ISO-15216 method and the bead-based method except for cereals (cornflakes) for which the bead-based assay had previously failed to recover FCV and MNV (10). Quantification of the recovered viral genomes was performed at different time points using ddRT-PCR and the results are presented as log reduction compared to T_0_. As shown in Figure 1, the reduction in viral load was less than 0.5 log during the 4-week incubation for all the studied viruses and the LMF commodities. This finding indicates that the genomes of FCV, MNV, and HAV remain stable on LMF during long-term incubation. Moreover, despite the premise that the employment of bead-based assays leads to extraction of “intact” viral capsids and therefore provide a better insight into the presence of infectious viral particles in food commodities, the results of the bead-based assay and the ISO-15216 methods demonstrate similar decline rate in viral genome copy number during the 4-week storage (Figure 1).

**Figure 1.**
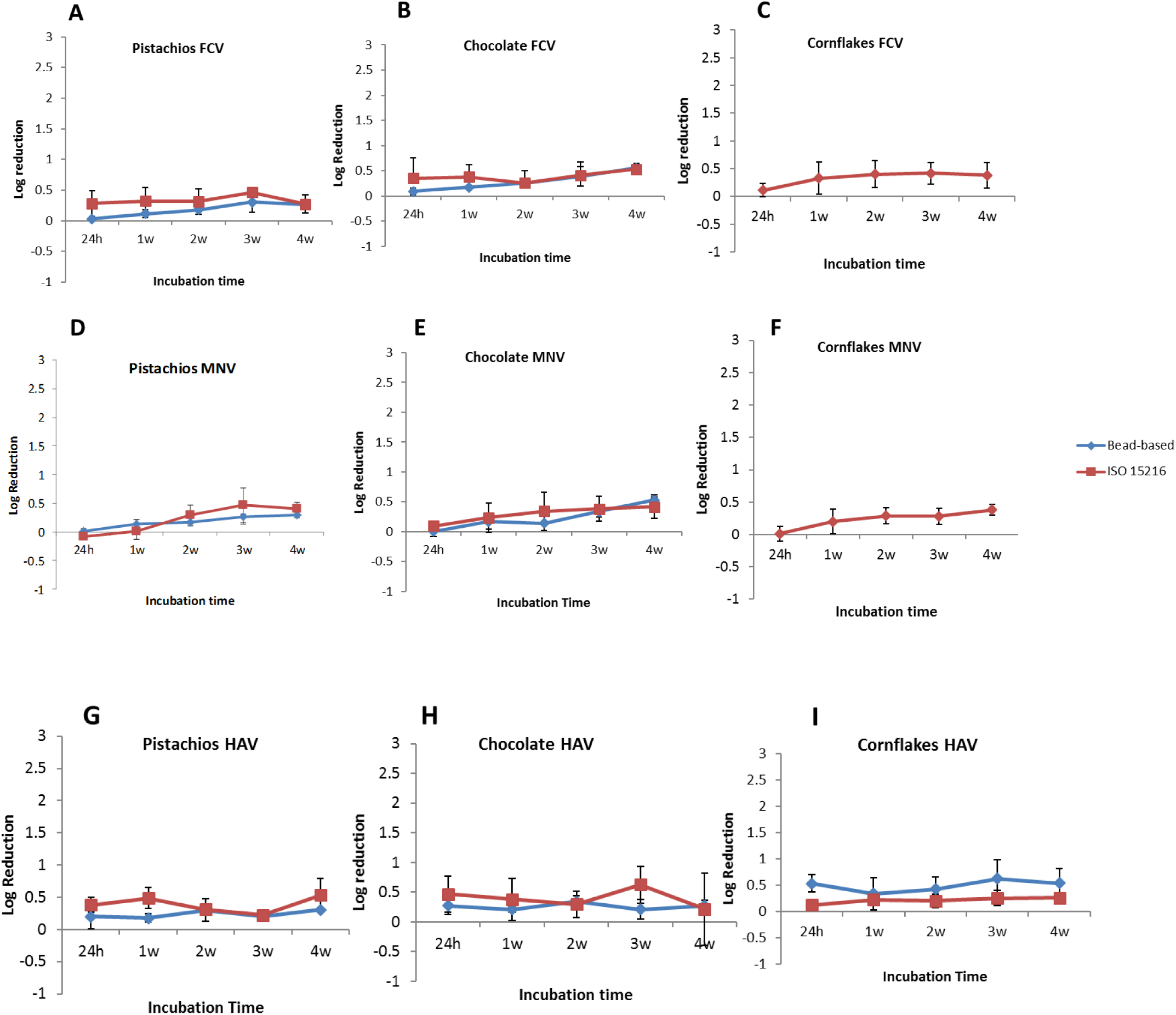
Log reduction of viral genome copies compared to T_0_. FCV A-C, MNV D-F, and HAV G-I recovered by the ISO-15216 method and the bead-based method in inoculated pistachios, chocolate, and cornflakes over 4-week incubation determined by ddRT-PCR. Error bars represent standard deviation.

To further examine the viral survival in LMFs, we performed infectivity assay (plaque assay) on viruses that were isolated at different time points post-inoculation and compared the infectivity results to T_0_. As shown in Figure 2, while the infectivity of FCV decreases gradually in all studied LMFs during the 4-week incubation time to less than 1 log (0.83 ± 0.1 log) compared to T_0_ (Figure 2A), the overall infectivity of MNV and HAV reduced to 1.2 ± 0.2 log and 1.34 ± 0.14 log (Figure 2B, and C), respectively. These data are in line with previous reports that foodborne viruses are persistent in foods and the environment over a long period of time.

**Figure 2.**
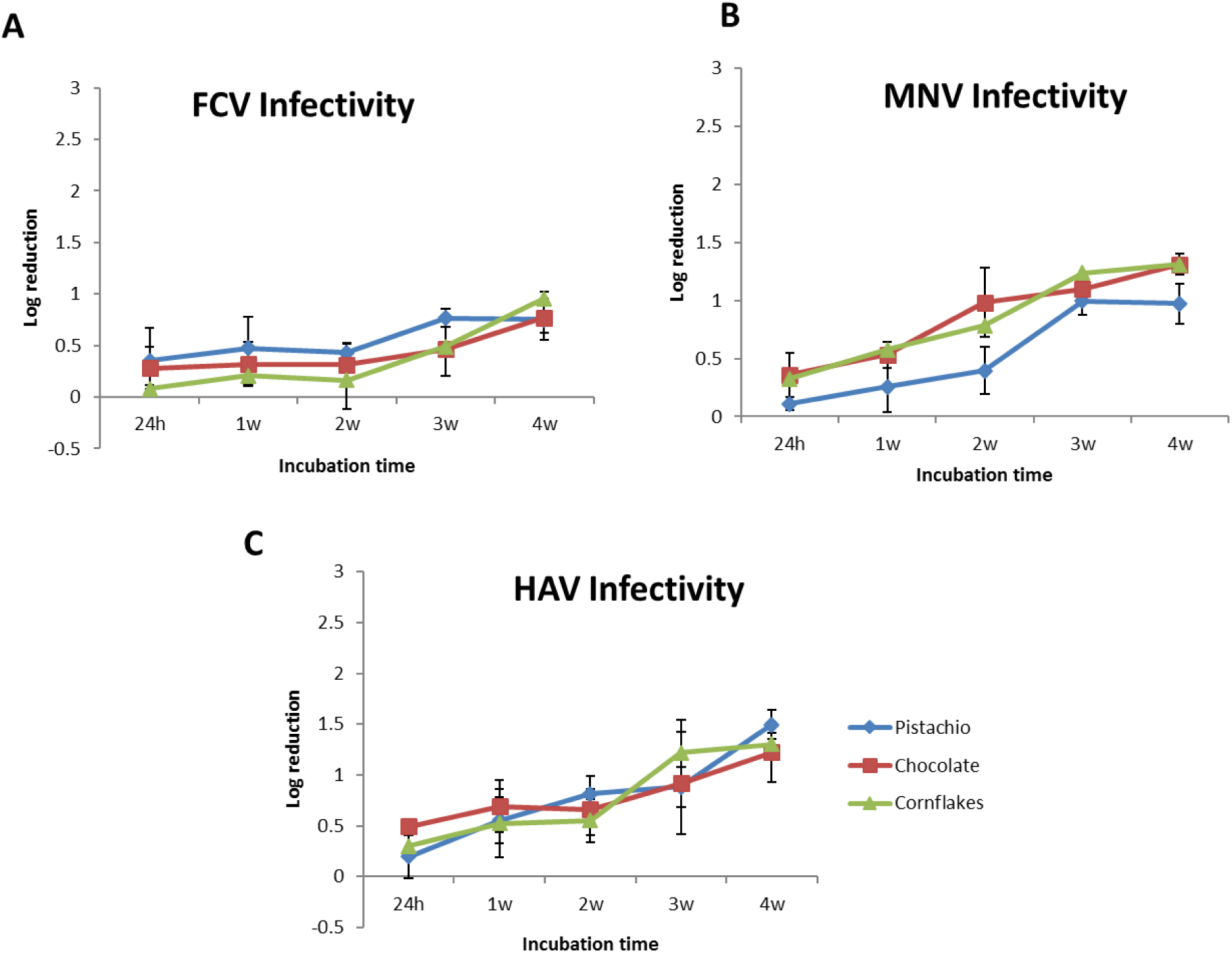
Log reduction in infectious viral particles for A) FCV, B) MNV, and C) HAV over 4-week incubation determined by plaque assay.

### Genomic analysis of the surviving viruses

To determine genomic changes that might have occurred during the incubation on LMF, or whether the incubation on LMF would select for certain quasispecies, we performed next-generation sequencing (NGS) on extracted viruses from T_0_ and week 4 (W4) samples to compare their genomes. For this purpose, we employed a metagenomic WGS method that we had successfully used for WGS analysis of human norovirus from clinical samples (24, 25). The metagenomic approach also allowed for capturing all the possible variations within the extracted population, and the *de novo* assembly further reduced any sequencing bias. Genome coverage varied from 14 to 2827 fold for the extracted viruses with an average of 436 fold (Table 2). While 10 single nucleotide variations (SNVs) were observed in FCV genome at W4, only 2 SNVs were identified in MNV and HAV genomes (Table 2) compared to their respective T_0_ consensus sequences. In addition, some of SNVs led to an amino acid change compared to the consensus input genome, whereas some of variants were synonymous (Table 2). While for some SNVs such as FCV-2195, and FCV-2436 the change is more eminent with the ratio of T811:A5 and T33:C0, respectively, other SNVs, like FCV-1358 (C12:T13), FCV-1834 (A107:G104), and MNV-4935 (C45:T49) the prevalence of the identified variants is close to the consensus nucleotide. This finding can further validate the notion that some variants exist at lower frequency, while others are commonly found within a viral quasispecies to the point that makes it difficult to identify the dominant variant.

**Table 2.** The list of single nucleotide variants (SNVs) that were found by NGS between the viruses isolated from W4 and T_0_. The position of each SNV including the open reading frame (ORF), the sequencing coverage for that position, the ratio between the identified SNVs, the food in which the SNVs were identified, and potential amino acid (aa) change were demonstrated.

| Virus | ORF | Position | Coverage | nt<br>Change | Ratio | Food | Time | aa Change |
| --- | --- | --- | --- | --- | --- | --- | --- | --- |
| FCV | 2 | 1265 | 38 | A-G | A: 30, G: 8 | Chocolate | T0-W4 | none |
| FCV | 2 | 1393 | 23 | T-G | T: 7, G: 16 | Chocolate | T0-W4 | none |
| FCV | 1 | 2195 | 818 | T-G | A: 5, C: 1, G: 1, T: 811 | Cornflakes | T0-W4 | I-S |
| FCV | 2 | 1262 | 2214 | A-G | A: 1682, C: 2, G: 519, T: 9 | Corn flakes | T0-W4 | E-G |
| FCV | 2 | 1277 | 2827 | T-C | A: 4, C: 1114, G: 3, T:<br>1706 | Cornflakes/Pistachios/Chocolate | T0-W4 | I-T |
| FCV | 1 | 2436 | 33 | T-C | T: 33, C: 0 | Pistachios | T0-W4 | none |
| FCV | 1 | 2826 | 18 | C-T | C: 16, T: 2 | Pistachios | T0-W4 | I-V |
| FCV | 1 | 3699 | 14 | A-G | A: 14, G: 0 | Pistachios | T0-W4 | T-A |
| FCV | 2 | 1328 | 25 | C-T | C: 12, T: 13 | Pistachios | T0-W4 | S-A |
| FCV | 2 | 1834 | 211 | A-G | A: 107, G: 104 | Pistachios | T0-W4 | N-G |
| HAV |  | 199 | 75 | A-C | A:20, C:55 | Pistachios | T0-W4 | none |
| HAV |  | 2498 | 46 | T-C | T:11, C:35 | Pistachios | T0-W4 | L-F |
| MNV | 1 | 4532 | 102 | T-C | T:68, C:34 | Chocolate | T0-W4 | none |
| MNV | 1 | 4935 | 94 | C-T | C:45, T:49 | Chocolate | T0-W4 | Q-R |

### Inactivation by advanced oxidative process

In this study, we evaluated the efficacy of using the Advanced Oxidative Process (AOP) technology as a new method to inactivate foodborne viral pathogens in LMFs. The AOP unit has 4 components; UV-C light, ozone, heat, and hydrogen peroxide vapor. The unit consisted of a UV light box that had a combination of UV-C (254 nm) and ozone generating (185nm) lamps. Hydrogen peroxide was introduced via a vaporizer at 6% concentration. A heating fan functioned to distribute the hydrogen peroxide, in addition to maintaining the temperature around 48°C (Figure 3).

**Figure 3.**
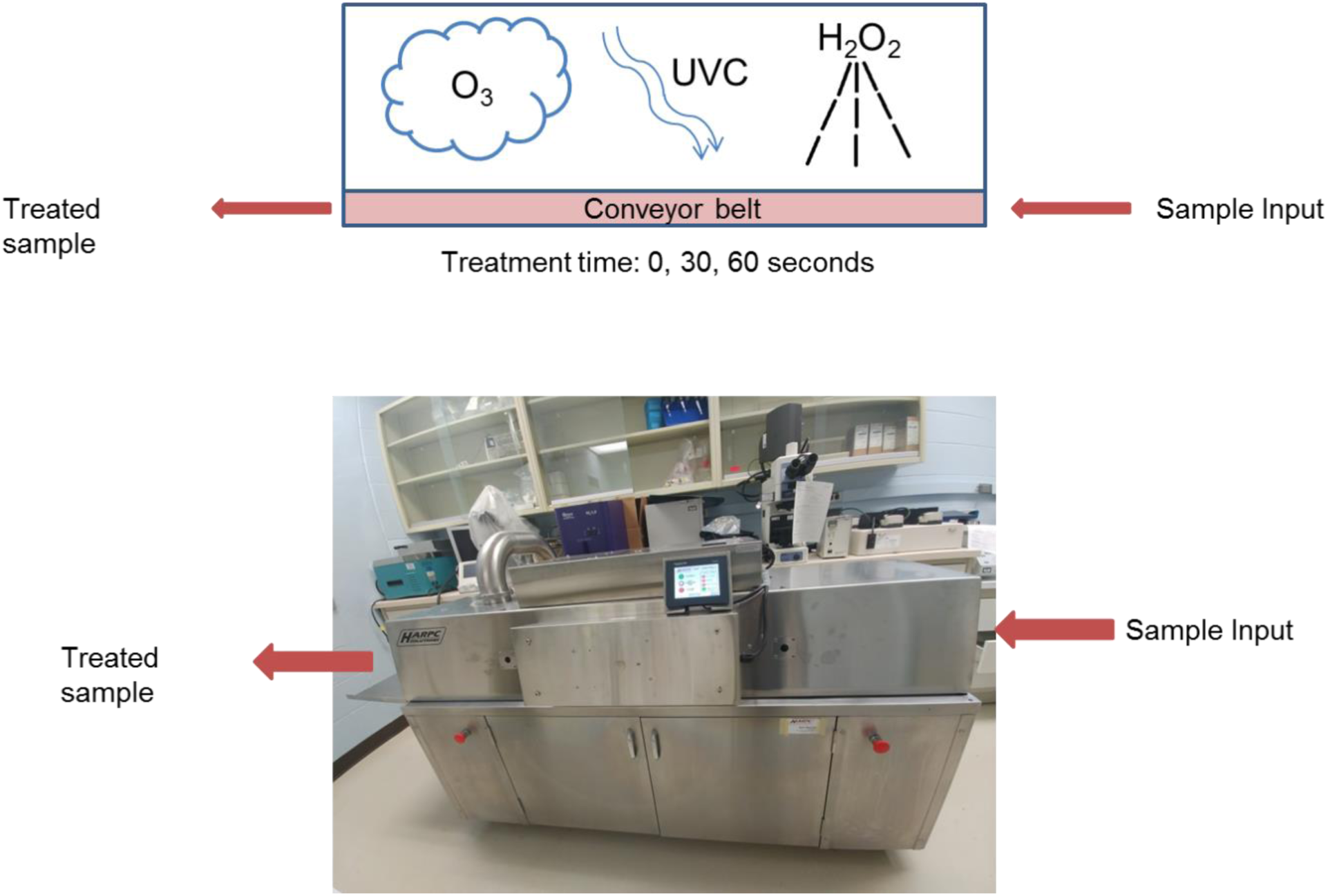
Inactivation of viruses on LMFs by Advanced Oxidative Process (AOP). Inoculated pistachios, chocolate, and cornflakes were treated with AOP for 0 minutes (untreated), 30 seconds, and 60 seconds. Virus recovery was done using the ISO-15216 method and quantified by ddRT-PCR and plaque assay.

Artificially inoculated pistachios, chocolate, and cornflakes with FCV, MNV and HAV were placed on a petri dish and kept in the AOP unit for 30 or 60 seconds. Viral inactivation was examined by comparing the viral titre of the recovered viruses from treated samples to the inoculated but untreated samples using both plaque assay and ddRT-PCR (Figure 4).

**Figure 4.**
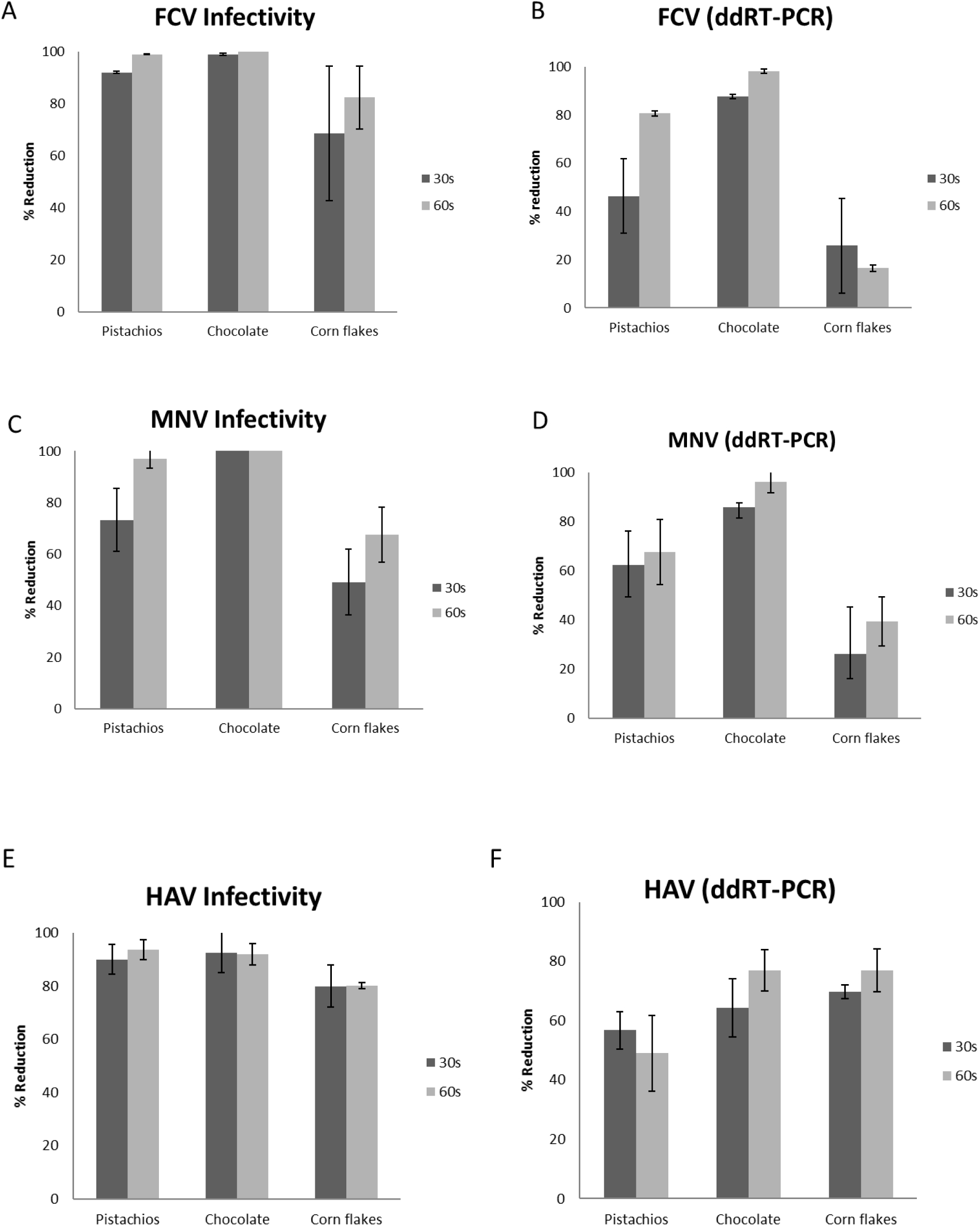
The percentage reduction after inactivation with AOP for 30 and 60 seconds for FCVA-B, MNV C-D, and HAV E-F in all three LMFs.

As shown in Figure 4A, FCV inactivation, examined by plaque assay, was close to 100% for chocolate, which has flat surface and about 95% and 75% for pistachios and cornflakes respectively. Also, the data suggest that the increase in the treatment time from 30s to 60s did not significantly enhance viral inactivation. However, the analysis of viral inactivation by examining the survival of the FCV genome by ddRT-PCR in Figure 4B reveals decreased inactivation compared to the infectivity assay (Figure 4A). This indicates that a proportion of surviving genomes that are quantified by ddRT-PCR do not belong to infectious particles.

Similar observations were made regarding the MNV inactivation (Figure 4C-D); no infectious viruses were recovered from the treated chocolate (30s and 60s), indicating that viral inactivation was 100%, while the inactivation in pistachios and cereals was approximately 97% and 68% respectively for 60s treatment. Like FCV, the MNV genome was more resistant to inactivation by AOP (Figure 4D) and demonstrated reduced inactivation compared to MNV infectivity assay (Figure 4C), suggesting that all recovered genomes might not be infectious.

Finally, there is no statistically significant difference in inactivation of HAV in chocolate and pistachios (93% and 92%, respectively), however, inactivation of HAV in cornflakes was only 80% (Figure 4E). Similar to observations made for FCV and MNV, the genome of HAV demonstrated a reduced inactivation by AOP treatment and the duration of treatment did not have a significant effect on inactivation (Figure 4F). Altogether, these data suggest that the viricidal effect of AOP treatment depends on the matrix and the surface of the treated LMF.

## Discussion

We have formerly compared the efficacy of the ISO 15216:2017 method with the magnetic beads assays for the isolation of HAV, MNV and FCV in shelled pistachios, chocolate liqueur and cereals and demonstrated that higher recovery rates can be obtained by the bead-based assays compared with the ISO-15216 method (10). However, the bead-based method failed to recover FCV and MNV from cornflakes and for this reason, the bead-based assay was not employed to examine FCV and MNV persistence in cornflakes (10).

Another advantage of the ISO-15216 method over the bead-based assay is that it allows for performing infectivity assay on the precipitated viruses. To date, we have not found an efficient method for the elution of the captured viruses by magnetic beads without having a drastic negative impact on their infectivity. Therefore, the viruses that were captured by the bead-based method were not subjected to plaque assay.

The infectivity survival data revealed that FCV is slightly more persistent in LMF compared to MNV and HAV, as the overall reduction in infectivity at W4 was less than 1 log for FCV, while the loss in infectivity of MNV and HAV was around 1.2 log and 1.34 log, respectively. Our findings are in accordance with the previous reports about survival of foodborne viruses on food-contact surfaces (30).

Except for cornflakes, the persistence of the viral genome was examined using two different extraction methods; the bead-based method and the ISO 15216 method. Both methods generated similar results with a reduction of approximately 0.5 log for all three viruses, with similar pattern of viral genome reduction rate over the incubation time in all tested LMFs (Figure 1). The reduction of the viral genome is significantly lower than the decrease in viral infectivity, which suggests that the presence of the viral genome does not necessarily correlate with the presence of infectious particles. Thus, the bead-based assay might not be a true indicator of viral infectivity. Altogether, these data demonstrate that these viruses persist in an infective state, regardless of the food matrix, for a relatively long period of time in LMFs and support the previous finding that low humidity might even be beneficial for viral persistence (31).

Even though viruses do not replicate in food, we were interested in examining whether the viruses that remain infectious after long incubation on LMFs, are genetically different from the viruses that did not go through the same process. In other words, we set out to examine the potential genetic alterations that might render the virus higher chance of survival in LMFs. For this purpose, we employed a bias-free metagenomic approach for deep-sequencing of the viruses (FCV, MNV, and HAV) recovered from T_0_ and W4. Bias was further reduced by using *de novo* assembly. The coverage-depth made it possible to identify all variants at each nucleotide position where an SNV was detected compared to the consensus sequence of T_0_. Only two SNVs were found for W4 sequences of HAV and MNV, while the number of detected SNVs were considerably higher for FCV. Whether the higher genetic diversity within the surviving FCV quasispecies leads to better chance of long-term survival, needs to be further investigated.

It has been estimated that class I recall of LMFs due to pathogen contamination, leads to a median loss in corporate value of $243,000,000 in the United States (18). Therefore, strategies that could reduce the risk of pathogen contamination, not only reduce the healthcare and societal costs associated with foodborne illnesses, but also have positive economic impact on the industry. Thus, novel and effective pathogen control technologies need to be developed, tested and implemented for improved food safety of LMFs. For this reason, we have collaborated with industry to examine the efficacy of the AOP system against foodborne viral pathogens.

Although the effect of UV-C, ozone, and hydrogen peroxide for viral inactivation has been evaluated individually, the combined viricidal efficacy of these strategies has never been examined. The AOP system enabled us to employ all these strategies in a single unit (Figure 3). We observed while the AOP treatment led to 100% (4 log) inactivation of FCV and MNV in chocolate (Figure 4), the treatment of inoculated pistachios and cornflakes caused less than 90% (1 log) reduction in viral infectivity. This finding indicates that either the food matrix or the surface of treatment can influence the outcome of inactivation. The effect of surface on viral inactivation has been shown previously for hydrogen peroxide vapor (32), ozone (33), and UV-C (34) individually. Also, it has been demonstrated that the presence of organic matter diminishes the ozone treatment efficacy (33). We speculate that the successful application of AOP relies on the treatment components i.e. UV-C, ozone and hydrogen peroxide vapor reaching all the virus particles directly and if the viruses are present in cracks, crevices or openings in the food and surfaces, the viruses may be protected and thus survive. Further AOP research works needs to be performed on a combination of food and surfaces to determine whether the surface of treatment or the matrix can affect the inactivation outcome.

In conclusion, we employed the bead-based method and the ISO 15216 method to demonstrate that foodborne viruses are persistent for a long period in LMFs. Furthermore, we demonstrated that depending on the food matrix and its surface, the AOP treatment could be an effective means to improve the virological food safety of LMFs.

## Acknowledgements

This study was in part supported by the grant from ILSI North America and the Microbiology Research Division of the Bureau of Microbial Hazards, Health Canada. ILSI North America is a public, non-profit science foundation that provides a forum to advance understanding of scientific issues related to the nutritional quality and safety of the food supply. ILSI North America receives support primarily from its industry membership. ILSI North America had no role in the design, analysis, interpretation, or presentation of the data and results. We would like to thank Dr. Jeffrey Farber (University of Guelph) for his insightful comments throughout the project as well as Dr. John Austin and Dr. Ana Pilar (Health Canada) for providing comments for the improvement of the manuscript.

